# Outlier detection for improved differential splicing quantification from RNA-Seq experiments with replicates

**DOI:** 10.1101/104059

**Authors:** Scott Norton, Jorge Vaquero-Garcia, Yoseph Barash

## Abstract

**Motivation:** A key component in many RNA-Seq based studies is contrasting multiple replicates from different experimental conditions. In this setup replicates play a key role as they allow to capture underlying biological variability inherent to the compared conditions, as well as experimental variability. However, what constitutes a “bad” replicate is not necessarily well defined. Consequently, researchers might discard valuable data or downstream analysis may be hampered by failed experiments.

**Results:** Here we develop a probability model to weigh a given RNA-Seq sample as a representative of an experimental condition when performing alternative splicing analysis. We demonstrate that this model detects outlier samples which are consistently and significantly different compared to other samples from the same condition. Moreover, we show that instead of discarding such samples the proposed weighting scheme can be used to downweight samples and specific splicing variations suspected as outliers, gaining statistical power. These weights can then be used for differential splicing (DS) analysis, where the resulting algorithm offers a generalization of the MAJIQ algorithm. Using both synthetic and real-life data we perform an extensive evaluation of the improved MAJIQ algorithm in different scenarios involving perturbed samples, mislabeled samples, no-signal groups, and different levels of coverage, showing it compares favorably to other tools. Overall, this work offers an outlier detection algorithm that can be combined with any splicing pipeline, a generalized and improved version of MAJIQ for differential splicing detection, and an evaluation pipeline researchers can use to evaluate which algorithm may work best for their needs.

**Availability:** Program is accessible via http://majiq.biociphers.org/norton_et_al_2017/

**Contact:** http://yosephb@upenn.edu

**Supplementary information:** Supplementary data are available at *Bioinformatics* online.

## 1 Introduction

Alternative splicing, the process by which segments of pre-mRNA can be arranged in different subsets to yield distinct mature transcripts, is a major contributor to transcriptome complexity. In humans, over 90% of multi-exon genes are alternatively spliced, and most of those exhibit splicing variations which are tissue- or condition-dependent [13]. This key role of alternative splicing (AS) in transcriptome complexity, combined with the fact that aberrant splicing is commonly associated with disease state [20], has led to various research effort. These include efforts to accurately map transcriptome complexity and identify splicing variations between different cellular conditions, across developmental stages, or between cohorts of patients and controls.

Detection of splicing variations and the mapping of transcriptome complexity has been greatly facilitated by the development of high-throughput technologies to sequence transcripts, collectively known as RNA-Seq. Briefly, RNA from the cells of interest, typically poly-A selected or ribo-depleted, are sheared to a specific size range, amplified, and sequenced. In most technologies used today, the resulting sequence reads are typically around 100bp with read number varying greatly, from around 20 to 200M reads. The shortness of the reads, their sparsity, and various experimental biases make inference about changes in RNA splicing a challenging computational problem [1]. Consequently, many studies include several RNA-Seq samples which are biological replicates of the conditions being studied. Replicates are a key component in helping researchers distinguish between the biological variability of interest (e.g. differences between two tissues) and variability associated with experimental or technical factors. However, what constitutes a “good” replicate or an outlier experiment is not always clear. Intuitively, in the context discussed here, an outlier is a sample which exhibits disproportionately large deviations in exon inclusion levels compared to other samples from the same experimental condition (biological replicates). An outlier could be the result of a failed experiment or of some hidden deviation in sample identity (i.e. different tissue source). Remarkably, despite the obvious importance of the question of what constitutes an outlier, this question has been mostly ignored in the literature. Instead, researchers are left to define outliers based on some heuristics which may carry unconscious biases or otherwise be less than ideal. For example, in a recent work focused on good practices for RNA-Seq data analysis, the authors recommend performing PCA analysis to see if samples from the same condition “cluster together” but admit “no clear standard exists for biological replicates” [5]. As we demostrate here, the presence of outliers can have deleterious effects on algorithms that aim to detect differential splicing (DS) between groups of replicates from different conditions. Thus an important contribution of this work is to formalize what constitutes an outlier replicate and suggest a method which researchers could use to detect outliers.

A second contribution of this work is in assessing the effect of outliers on DS algorithms and, more broadly, a general pipeline for assessing the performance of DS algorithms. Generally, algorithms that aim to quantify differential splicing from RNA-Seq can be divided into two classes. The first, which includes tools such as RSEM [10] and Cuffdiff [18], aims to quantify full gene isoforms, typically by assuming a known transcriptome and assigning the observed reads to the various gene isoforms in the given transcriptome database. The second class of algorithms, which includes rMATS [16] and DEXSeq [2], works at the exon level, detecting differential inclusion of exons or exonic fragments. Some algorithms, such as SUPPA [6], can be considered a hybrid as they collapse isoform abundance estimates from other algorithms (e.g. SailFish [15] or SALMON [14]) to compute relative exon inclusion levels. Previous works showed that for the task of differential splicing, quantification algorithms that work at the exon level generally perform better since they solve a simpler task and are less sensitive to isoform definitions or RNA-Seq biases within samples or along full isoforms [12]. Thus, for the comparative analysis section of this paper, we focus on the second class of algorithms, and specifically the commonly-used DEXSeq, rMATS, as well as the recently updated SUPPA, all of which support replicates.

Finally, a third contribution of this work is in adapting our recently published DS algorithm, MAJIQ [19], to automatically detect and downweight samples and splicing variations suspected to be outliers. MAJIQ, along with its matching visualization package VOILA, offers an effective method to detect, quantify, and visualize differential splicing between experimental conditions with or without replicates. Besides the details of its statistical model, two key features distinguish MAJIQ from the algorithms mentioned above. First, MAJIQ does not quantify whole gene isoforms as the first class of algorithms described, or only previously defined AS “types” (eg cassette exons), as the second class of algorithms. Instead, MAJIQ defines a more general concept of “local splicing variations”, or LSVs. Briefly, LSVs are defined as splits in a gene splice graph where a reference exon is spliced together with other segments downstream (single source LSV) or upstream of it. Importantly, the formulation of LSVs enables MAJIQ to capture all previously defined types of AS but also many other variations which are more complex. The second important distinguishing element of MAJIQ is that it allows users to supplement previous transcriptome annotation with reliably-detected *de-novo* junctions from RNA-Seq experiments. Despite its novel features and capabilities, the original MAJIQ algorithm was not built to handle outlier replicates. Indeed, we show here that such outliers can significantly affect MAJIQ’s performance. In contrast, the new generalization of MAJIQ that uses the weighting scheme we develop here, offers a DS algorithm which is not only robust to outliers but can also salvage useful information from outlier samples. In order to demonstrate this improved performance, we perform extensive comparative analysis of the new MAJIQ to other algorithms in terms of reproducibility of inferred differential splicing events, false positives when no biological signal is expected, and independent validation using RT-PCR at varying degrees of read coverage.

The rest of this paper is organized as follows: Section 2.1 formulates the model for outlier detection and how the resulting sample weights can be incorporated into the MAJIQ model. Section 2.2 then extends the global weight per-sample outlier model to a local, per-LSV model. Section 2.3 describes the methods used to evaluate algorithm performance and to generate synthetic data. Section 2.4 describes one method for artificially introducing an outlier in a group of replicates by perturbing splice inclusion levels for a subset of LSVs. Finally, Section 3 details the comparative analysis on synthetic and real data of several algorithms for detecting differential splicing using replicates, followed by a discussion and future directions.

## 2 Methods

### 2.1 Outlier weight model

Let *T* be the set of RNA-seq experiments presumed to be biological replicates for which alternative junction inclusion is to be measured, and let *t ∈ T* be one such experiment. All experiments constitute observations of reads mapping to *L* LSVs. Let *i* = 1, 2*, …, L*, and let *J* be the number of junctions in LSV *i* with indices *j* = 1*, …, J*. Then 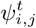 is the inclusion ratio of junction *j* of LSV *i* within experiment *t*, such that 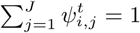. Importantly though, we never observe 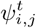 directly, but rather junction spanning reads drawn according to it. Consequently, the MAJIQ model does not infer 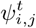 as a point estimate but rather as a posterior beta distribution, denoted 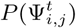 or, with slight abuse of notation, simply 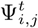 [19]. Another important point is that MAJIQ assumes that all the experiments in *T* are replicates of the same biological condition (tissue type, treatment, disease state, etc.) such that they generally share the underlying (hidden) junction inclusion levels, denoted 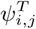, ∀*i, j*. Under this model, an *outlier s ∈ T* is an experiment which does not represent the same experimental or biological condition as the majority of the experiments in *T* such that 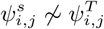 for a disproportionately large set of LSVs.

In order to make the above definition formal we need to specify a way to estimate 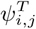, a measure of disagreement between 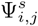 and 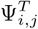, and a measure for how much we expect 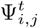 to deviate from 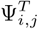 under normal conditions i.e. when *t ∈ T*. Intuitively, if an experiment agrees on inclusion levels with the majority of the group, its individual 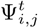 posterior distribution should be similar to the median of the group, which we use as a proxy for 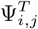. In contrast, the 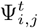 distribution for a group outlier should have high divergence from the median. We quantify the degree to which an experiment’s estimate of Ψ resembles the median using *L*1 divergence, defined as

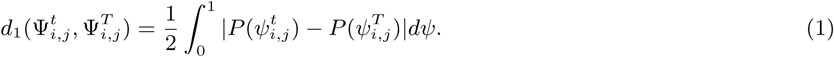

Using the above definition we then define the deviation of sample *t* for LSV *i* as 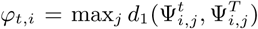. Under normal conditions each experiment *t* should have a similar distribution of *φ*_*t,i*_*, ∀i*, but for outliers this distribution will be skewed towards higher values, reflecting an accumulation of LSVs for which *φ*_*t,i*_ is high. This skew towards high *φ*_*t,i*_ values is illustrated in Figure 1. To capture such skewness, we find the number *K*_*t*_ = *|*{*i* : *φ*_*t,i*_ *> τ*}| of highly-disagreeing LSVs for each replicate, where *τ* is an arbitrary threshold for strong disagreement. In the experiments described below we used *τ* = 0.75 but we find that results were robust to changes in *τ* value as long as it was reasonably high (0.5 *< τ <* 0.9, data not shown). Theoretically, if all experiments are equally “well-behaved” we expect *K*_*t*_ to follow a binomial distribution *K*_*t*_ ∼ Binom(*K*_*T*_*, p*), where 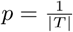. In practice, we find the distribution of *K*_t_ to have higher dispersion which we model using a Beta Binomial such that 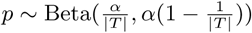. Setting *α* = 15.0 offered robust results in all the experiments and datasets we tested (data not shown). Finally, we set the weight *ρ*_*t*_ ∈ [0, 1] representing sample *t* belongs to group *T* using following ratio test: We compute the probability mass for the tail of the beta binomial probability *P*_*BB*_ for the observed number of highly-disagreeing LSVs *K*_*t*_, compared to the excepted 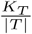 under the null-hypothesis expectation of equal sharing:

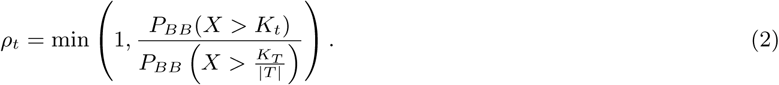

**Figure 1:**
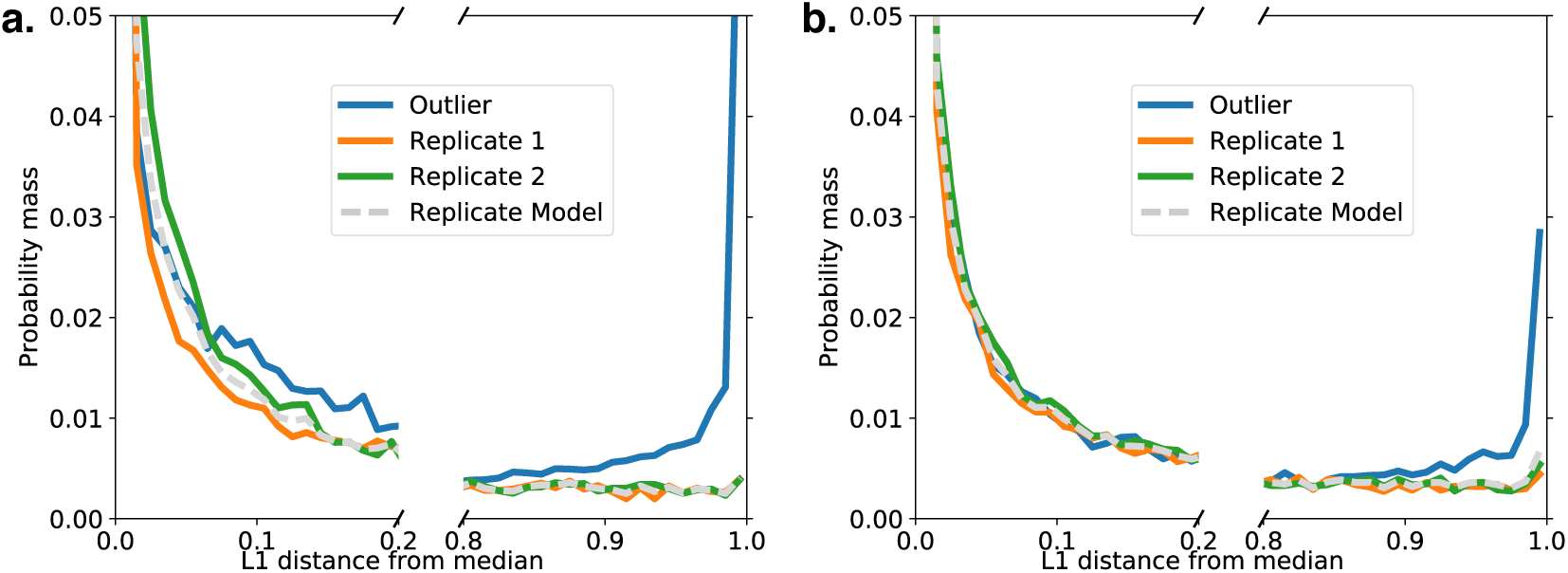
Per-experiment empirical probability distribution over LSVs *L*_1_ divergences *φ*_*t,i*_ as defined in the main text. Each solid line represents a different RNA-seq experiment (*t*), while the gray dashed line represents a weighted average based on *ρ*_*t*_. **(a)** A swap case of three “replicates” from [22] with two liver samples (green, red) and an outlier, “mislabeled”, muscle sample (blue). **(b)** Same as in (a) but a less extreme case where less LSVs have extreme divergence from the median. The blue outlier was produced by synthetically perturbing 10% of the LSVs by 10% in a cerebellum sample at the same coverage level (*γ* = 1.00, see Section 2.4). In both scenarios most LSVs have low divergence (*φ*_*t,i*_ *<* 0.2) but a small fraction of LSVs in the outlier have *φ*_*t,i*_ *>* 0.5, resulting in outlier weight dropping towards zero (*ρ*_*t*_ *<* 10^*−*3^). See Figure S1 for the distribution of *φ*_*t,i*_ when there is no outlier.

In the new MAJIQ implementation users can now invoke the option to compute these weights for groups of replicates to quickly detect possible outliers. We note that since the LSVs in the experiments are expected to be quantified in any case, the computational overhead associated with these weight computation is negligible.

In addition, the above global weight per-experiment *ρ*_*t*_ can be plugged into MAJIQ to produce a generalized version of the previous MAJIQ algorithm:

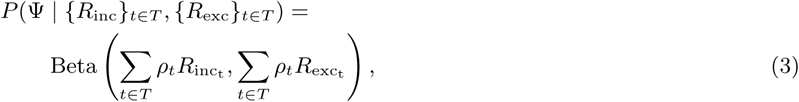

where *R*_inc_*, R*_exc_ represent estimated read rates supporting inclusion or exclusion of a specific junction. Details of how these read rates are derived from observed reads are given in [19]. In the evaluations that follow we refer to the resulting MAJIQ model with global weights per experiments as MAJIQ-gw while the previous MAJIQ model with no weighting scheme is denoted MAJIQ-nw.

### 2.2 Local weights

The previous section described MAJIQ-gw as a way to globally weigh experiments using *ρ*_*t*_ so that outliers can be effectively detected and suppressed. However, by doing this we run the risk of discarding valuable information: Even if a sample is a *bone-fide* outlier, it is quite likely that this is a result of a subset of LSVs, while others (typically the majority, as depicted in Figure 1) of the LSVs are good representatives of the condition of interest *T*. This phenomenon is nicely captured in Figure 1. To balance the benefits of outlier removal with the possible loss in statistical power we develop a local weight model (*ν*_*t,i*_ ∈ [0, 1], *∀t, i*) based on the distribution of *φ*_*t,i*_ we expect to find in “well-behaved” replicates.

First, we estimate the null distribution of “well-behaved” replicates by weighting the per-experiment densities of *φ*_*t,i*_ by *ρ*_*t*_. In Figure 1, this distribution is represented by the light gray line. Next, for each *t, i*, we estimate the matching LSV’s L1 divergence falling within a neighborhood of *φ*_*t,i*_ under the distribution for experiment *t* compared to the null model. Explicitly, we define

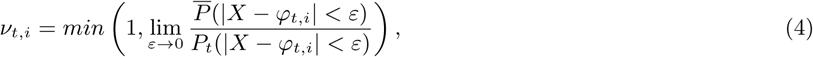

where 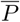 is the null distribution of *φ*_*t,i*_ and *P*_*t*_ is the distribution of *φ*_*t,i*_ for experiment *t*. Plugging *ν*_*t,i*_ into Equation 3 instead of *ρ*_*t*_ gives us the local weight model, which we denote MAJIQ-lw.

### 2.3 Performance evaluation metrics

There is an inherent challenge in assessing the accuracy of methods for RNA-Seq analysis since the underlying true values are rarely known. Some works use synthetically-generated samples with specific transcripts spiked at different concentrations which may be very different from real life samples, while others resort to synthetic sequencing data generation under various simplifying assumptions. Instead, we focus here on using real life data with multiple replicates to assess *reproducibility* in different experimental setups as a mean of assessing the performance of algorithms that quantify Ψ or differential splicing (DS). Here DS is measured as inclusion changes between two conditions *T*_1_, *T*_2_ such that 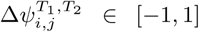, where the matching posterior distribution given observed reads is denoted 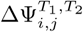. Specifically, we use a reproducibility measure (*RR*) similar to the irreproducible discovery rate (IDR), which has been used extensively to evaluate ChIP-Seq peak calling methods [11] and, more recently, for methods detecting cancer driver mutations [17]. Conceptually, *RR* is a rank-based statistic, agnostic of an algorithm’s model or scoring metric. It measures the proportion of high-ranked events (e.g. ChIP-Seq peaks or differentially-spliced events) that are also observed in a second, independent iteration of the same experimental procedure using biological replicates. To compute the *RR*, an algorithm *A* is run on a “training” set, denoted *S*1, and outputs the number of differentially-spliced events (*N*_*A*_), ranked by their relative significance or score. For any *n ≤ N*_*A*_, we then compute the size of the subset of events(*R*_*A*_(*n*) = *n*^*’*^ *≤ n*) of those *n* events which are ranked in the *n* highest ranking events in a second “hidden” test set (*S*2). In [19] the reproducibility graphs plotted 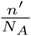 against 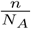 with perfect reproducibility corresponding to a 45^*°*^ line. This setup was unsuitable for the algorithms evaluated here as these varied greatly in their estimated *N*_*A*_ (see Section 3). We therefore opted to plot 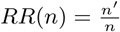 against *n* which captures the reproducibility ratio for any given *n* and is not sensitive to the definition of *N*_*A*_. See Figure S2 for an illustration of how the previous and current reproducibility plot definition relate to each other.

While useful, the *RR*_*A*_ and *N*_*A*_ statistics for a specific algorithm *A* may not be sufficient to assess an algorithm’s performance as they suffer from several caveats. First, both *RR*_*A*_ and *N*_*A*_ are not inherent characteristics of an algorithm *A* but rather a combination of an algorithm and a dataset. Furthermore, different algorithms may use different statistical criteria to call a splicing variation significantly changing. Consequently, their *N*_*A*_ may vary greatly. Second, reproducibility by itself is not a measure of accuracy as algorithms can be highly reproducible yet maintain a strong bias. Therefore in order to better assess accuracy of methods for differential splicing quantification, we perform two additional tests of performance. First, we assess a lower bound on the number of false positives (FP) by comparing sub-groups of samples from the same experimental condition. Consequently, both sub-groups being compared are expected to contain only similar within group variability. Thus, the significantly changing events between these two sub-groups (*N* ^*ns*^) are expected to be FPs with respect to the between conditions signals. However, since we cannot rule out inherent unknown bias even within the no-signal groups, we compute *R*(*N* ^*ns*^), expecting it to be close to 0. We then compute a conservative lower bound estimate on the False Discovery Rate (FDR) for a given algorithm *A* on dataset *D* as:

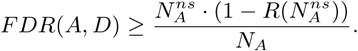

Finally, as a third measure for an algorithm’s accuracy, we used RT-PCR triplicate experiments from previous studies [19]. This measure is limited by the total number of events quantified, possible selection biases, and limitations of the experimental procedure. For example, for accurate quantification to be valid, experiments need to be executed in triplicates with careful reading of the gel bands. In practice, RT-PCR is sometimes used more qualitatively for calling changes, making reuse of published experiments less suitable for nuanced quantitative comparison. Also, PCR primers need to be carefully mapped to each method’s quantified event for valid comparisons, and past studies have mostly focused on classical AS events types rather than the more complex ones [19]. Nonetheless, carefully executed RT-PCR provide valuable experimental validation and is considered the gold standard in the field.

### 2.4 Synthetic perturbation

To observe the impact of disagreement on Ψ in a controlled fashion, we use a real replicate and perturb it to create a synthetic new pseudo-replicate outlier. Three parameters were used to control the sample’s perturbation: *θ ∈* [0, 1] determined the fraction of LSVs randomly selected to be perturbed, *δ ∈* [0, 1] controlled the shift of *E*[Ψ], and *δ >*= 0 set the relative coverage level in the perturbed sample. Specifically, we used the following procedure:

1. Set *θ ∈* [0, 1], *δ ∈* [0, 1], and *γ >* 0.
2. Randomly sample *L ⊂* LSVs with *|L|* = *θ|*LSVs*|*.
3. For *l ∈ L* with per-junction read rates *μ*_*l,j*_*, j* = 1*, …, J*:
  a. Estimate *E*[Ψ_*l,j*_] for each junction.
  b. Sample *ε ∼ U* (0, 1) and let

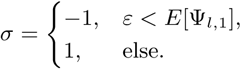
  c. Set 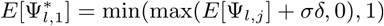.
  d. For 2 *≤ j ≤ J*, set

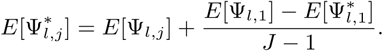
  e. For 1 *≤ j ≤ J*, set

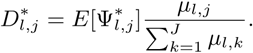
4. For *l ∈* LSVs *\ L*, set 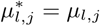.
5. For *l ∈* LSVs, set 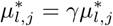 .

Observe that when *γ* = 1 and 0 ∈ {*θ, δ*}, the synthetic perturbation does not alter *μ* for any LSV. We measure the effect of variations in *θ*, *δ*, and *γ* on *ρ*_*T*_ and *RR* by applying the above Ψ perturbation to one replicate in set *S*_1_.

### 2.5 Mislabeled sample

In an extreme case, we explore the effects of mislabeling a sample. We simulate this by swapping out one replicate in the set *S*1 with a sample from a different condition within the same dataset.

### 2.6 Source data

In order to perform the type of evaluations described above, datasets with at least six replicates are required. The results described here were derived using RNA-seq experiments from a recent study by [22], which includes samples from twelve mouse tissues (heart, hypothalamus, adrenal, brown and white fat, cerebellum (Cer), liver (Liv), muscle (Mus), lung, aorta, brainstem, and kidney) across four circadian time points, repeated twice, with 60-120 million reads per experiment.

## 3 Results

To demonstrate the robustness of both the global (MAJIQ-gw) and local (MAJIQ-lw) weighting algorithms, we applied the synthetic perturbation procedure in Section 2.4 to one experiment in a group of three cerebellum samples from [22], varying each of the three hyper-parameters *θ, δ, γ* while holding the other two fixed. We compared these new generalizations of MAIJIQ to the previous algorithm (MAJIQ-nw) run with the outlier sample and to a case where we assume some heuristic (e.g. PCA) was able to detect and remove the outlier before the old algorithm was run (MAJIQ-rm). Figure 2 shows the effect of varying each hyper-parameter on the global weight associated with that experiment (log_10_ *ρ*_*T*_, left column), the number (*N*_*A*_, middle column) of LSVs reported as differentially spliced with high confidence (*P* (*|*ΔΨ*| >* 0.2) *>* 0.95%), and the reproducibility ratio (*RR*_*A*_(*N*_*A*_), right column). At *δ* = 0.6*, γ* = 1 (a-c, top row), the outlier’s weight *ρ*_*t*_ scales linearly in log scale to the fraction of LSVs perturbed (*θ*) such that perturbing 10% is sufficient to drop *ρ*_*t*_ below 0.1. Consequently, the original MAJIQ-nw detects up to approximately 900 false positives and reproducibility drops down to approximately 60%, while the *N*_*A*_ and *RR* for both weighting algorithms (MAJIQ-gw and MAJIQ-lw) remain stable (Figure 2b,c). Notably, the power of MAJIQ-lw to detect changing LSVs under control conditions of no outliers (*θ* = 0.0) is approximately that of the old MAJIQ-nw and slightly exceeds that of MAJIQ-gw, probably due to the heterogeneous nature of the replicates (different circadian time points). When outlier LSVs are added (*θ >* 0.0) MAJIQ-gw and even more so MAJIQ-lw maintain robust *RR* while being able to identify more changing LSVs (*N*_*A*_) compared to simply removing the sample as in MAJIQ-rm.

**Figure 2:**
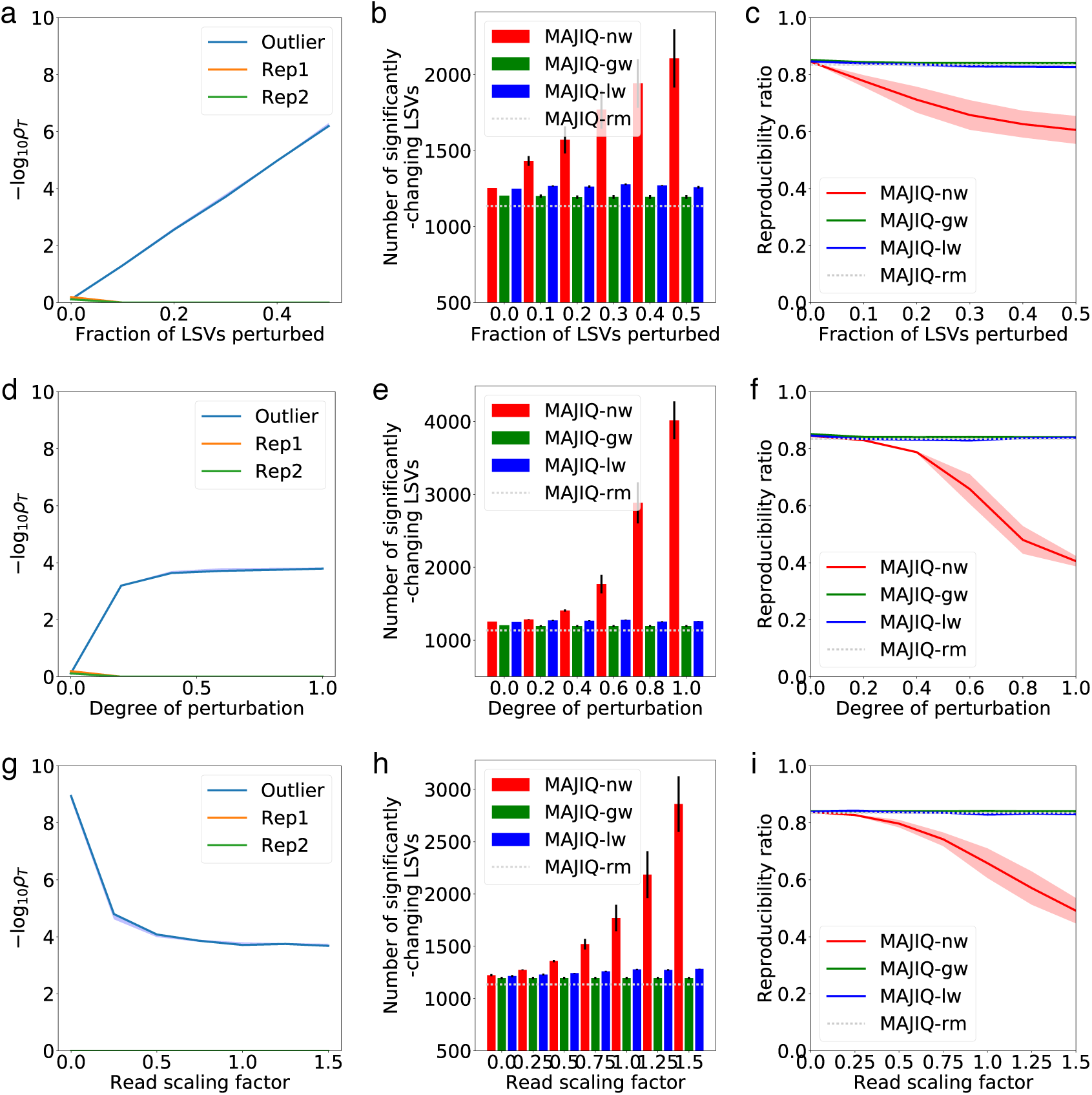
Synthetic perturbation of tissue replicates following the procedure in Section 2.4. In each test, three cerebella were compared against three Livers from [22], but one of the cerebella was perturbed using the given parameters. MAJIQ-nw is the previous algorithm equivalent to fixed weights (*ρ*_*t*_ = 1). MAJIQ-rm is a control case where we assume some heuristic (e.g. PCA) was able to detect the outlier and remove it before executing the previous fixed-weights MAJIQ. **a,d,g.** Effect on *ρ*_*t*_ for the perturbed “outlier” (blue) and unperturbed replicates (Rep1,2 in green and orange). **b,e,h.** Effect on the number *N*_*A*_ of events detected to have *P* (ΔΨ *>* 0.20) *>* 0.95 between the cerebellum and liver samples. **c,f,i.** Effect on the reproducibility ratio *RR*(*N*), defined in Section 2.3. **a-c.** *θ*, the fraction of LSVs perturbed, is varied between 0 and 0.5. At 0, the effect is the same as having no perturbation at all. **d-f.** *δ*, the maximal amount by which Ψ is perturbed, is varied between 0 and 1. *δ* = 0 is equivalent to no perturbation. **g-i.** *γ*, the “read scaling factor”, is varied between 0 and 1.5. When this factor is 0, it is functionally equivalent to a global weight of 0.

At *θ* = 0.3*, γ* = 1 (Figure 2d-f, middle row), increasing *δ* initially causes the weight on the outlier to decrease towards a positive infimum. For larger *δ >* 0.5, the original *N*_*MAJIQ−nw*_ increases over 3-fold with a corresponding 45% drop in *RR*_*M AJIQ*_. At *θ* = 0.3*, δ* = 0.6 (Figure 2g-i, bottom row), decreasing *γ* towards 0 causes the outlier weight to shrink below 10^*−*8^, suggesting that the algorithm is sensitive to a sample with very low read counts. Indeed, with very few reads the estimated Ψ distribution does not vary significantly from the prior in most cases but the sample has minimal effect on the posterior for the group as it contributes only few reads. This is reflected in the fact that the original MAJIQ-nw is not affected too at such low coverage levels. Unsurprisingly, increasing the read rates to 150% (Figure 2g-i for x-axis *δ* = 1.5) does not significantly affect the weight, which remains very low. It does, however, increase the unreliability of the original MAJIQ, more than doubling *N*MAJIQ-nw while nearly halving *RR*_MAJIQ-nw_. In all these cases, both MAJIQ-gw and MAJIQ-lw remain resistant to the perturbations and detect more changing LSVs than MAJIQ-rm.

Next, we evaluated the reproducibility with and without a sample swap for a large set of algorithms using the same dataset. In all cases, we used 3 cerebellum vs. 3 liver for the validation set 2. Set 1 included either 3 cerebellum compared with 3 liver samples (control, no swap) or two cerebellum and one muscle compared with three livers (swap case). Since the original dataset included circadian timepoints we made sure to match those for both sets. We chose to evaluate these tissues specifically because of the large number of reproducibly-changing LSVs between them [19], though similar results were observed for other tissues as well as for a different dataset from [9] (data not shown). To produce the reproducibility curves, we followed the following procedure. For the MAJIQ methods, *N*_*A*_ was defined over set 1 as the set for which *P* (*|*ΔΨ*| ≥* 0.2) *>* 0.95 as in [19]. Similarly, for SUPPA and rMATS we define *N*_*A*_ as the set of significant events as those with p-value *≤* 0.05, ranking events that were found significantly changing by ΔΨ and using a cutoff of ΔΨ *≥* 0.2. For DEXSeq we used the same significance cutoff of adjusted p-value *≤* 0.05 but since it only returns a log_2_ fold change and not ΔΨ we ranked events by this statistic with set a cutoff of log_2_ fold change *>* 4. We also tried ranking hits by FDR or p-value for these three methods but reproducibility was greatly degraded (data not shown). Finally, given the results described in the previous section, we only included MAJIQ-lw in this evaluation, and compared it against the previous MAJIQ-nw.

Figure 3a summarizes the results for the swap outlier evaluation procedure described in Section 2.5. Here, for each algorithm we also include a matching “Control” execution in which the swap did not occur. Also, given the results from the synthetic perturbations described above we focused here only on MAJIQ-lw compared to the previous MAJIQ-nw. One clear observation is the huge variation in the number of events reported as significantly changing (*N*_*A*_) by the different methods even when no outlier was present. This ranged from almost 30,000 for DEXSeq without outliers to 762 for MAJIQ-nw when a swap in performed. The reproducibility measures *RR*(*n*), which is agnostic of *N*_*A*_, was typically noisy for low *n*, which is to be expected. Nonetheless, reproducibility varied greatly between algorithms and for cases w/wo an outlier. MAJIQ consistently exhibited higher reproducibility compared to other algorithms regardless of *n*, with MAJIQ-lw and MAJIQ-nw practically tied in the control case with no outlier (*RR*(*N*_*A*_) = 84%). However, when an outlier was introduced the reproducibility of the previous MAJIQ-nw dropped to 77% while MAJIQ-lw identified more differential LSVs (777 vs 762) with higher reproducibility (82% vs. 77%). Interestingly, when introducing an outlier, SUPPA’s number of differentially spliced events dropped significantly (1857 vs 916), but unlike other algorithms its reproducibility in the range of *n ∈* [500, 1000] actually improved, increasing from 73% to 82% for *RR*(*N*_*A*_). As we show below, we suspect this counterintuitive improvement in reproducibility upon the introduction of an outlier is the result of high false positives with normal replicates, which the introduction of outliers may have helped to clear out. In addition, SUPPA’s output ran out of LSVs which could be mapped between Set1 and Set2 (marked with a bold circle in Figure 3a. This is likely the result of using only RefSeq annotated transcripts, as recommended in [6] to improve accuracy. In line with [16] we found rMATS not to be sensitive to outliers, though reproducibility was generally lower than MAJIQ and SUPPA.

**Figure 3:**
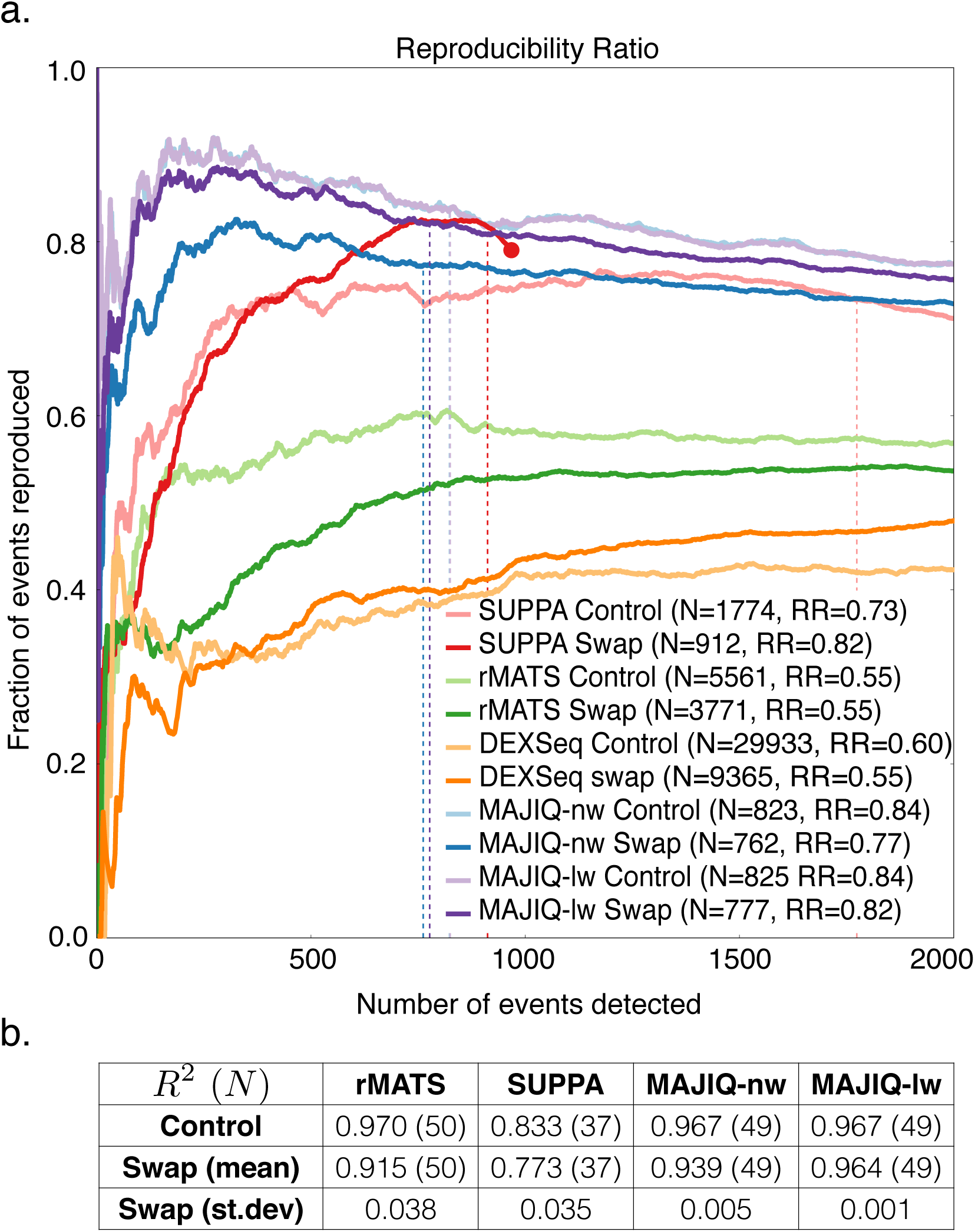
**(a)** Reproducibility plots for detection of differentially-spliced events between cerebellum and liver, with (Swap, solid line) or without (Control, faded line) a mislabeled muscle. Dashed vertical line denote the point in the graph matching the number of events reported as significantly changing by each method. Numbers in the legends represent this number (*N*_*A*_) along with the matching reproducibility value on the y-axis (*RR*(*N*_*A*_)). Bold circles mark cases where no more events were reproduced (see main text). Because DEXSeq and rMATS report more than 3000 significantly-changing transcripts, we present their extended RR curve in Figure S4. **(b)** The same control and swap experimental setup with accuracy assessed using 50 RT-PCR experiments from [19]. Values represent fraction of variations explained (*R*_2_) and the number of events detected in parentheses. DEXSeq is not included here since it does not output Ψ values. Scatterplots are presented in Figure S3. Mean and standard deviation estimates for the swap case are based on repeating the procedure three times, one for each cerebellum timepoint swapped out.

Next, we repeated the swap outlier experiment and measured its effect using 50 RT-PCR experiments from [19]. Figure 3(b) summarizes the accuracy of the various algorithms measured as the fraction of variations explained (*R*^2^). We found MAJIQ-lw, MAJIQ-nw, and rMATS gave similar *R*^2^ values in both the control and swap case, with MAJIQ-lw performing the best in the swap case. SUPPA exhibited lower *R*^2^ values, and detected fewer events. SUPPA’s lower number of events detected is likely due to its usage of Refseq only transcripts as noted above, but may also reflect selection bias as [19] originally selected events detectable by both rMATS and MAJIQ in order to compare these methods.

In order to assess the fraction of false positives reported by each method (FDR), we created “no-signal” groups from the same set of tissues used in Figure 3. Since the original dataset of [22] contained four circadian time points repeated twice for each tissue we created the no-signal groups from two biological samples from the same tissue (liver or cerebellum) compared to two other matching time points, 24 hours apart, in the same tissue. Since only two timepoints were used per group, MAJIQ-lw is equivalent to MAJIQ-nw in this case. The idea behind this experimental setup is to assess how many differentially spliced events a method reports between biological replicates of matched tissue and timepoints, and contrast that with the number of events reported as differentially spliced between different tissues and matched timepoints. Figure 4(a), shows MAJIQ reported a significantly smaller number of events in this test. Its estimated FDR, using the _*A*_ average of these as 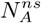 (See Section 2.3), was significantly lower and comparable to its *P* (*|*ΔΨ*| >* 0.2) *>* 0.95 statistic used to call significantly changing LSVs. By contrast, both DEXSeq and SUPPA have high false discovery rates (¿20%), with DEXSeq detecting over four thousand events under no-signal conditions.

**Table 1:** **(a)** *RR*_*A*_ and *N*_*A*_ for subsampling of cerebellum and liver samples from [22] to 100% (80 million reads), 50% (40 million reads), and 25% (20 million reads) of their original coverage, as reported by DEXSeq, rMATS, SUPPA, MAJIQ-nw, MAJIQ-lw, and MAJIQ-lw with the relaxed filter as explained in the text. **(b)** *R*^2^ values for accuracy of each method’s PSI estimation compared to RT-PCR quantifications under different levels of subsamplings. DEXSeq not included as it does not report Ψ values. Scatterplots matching these *R*^2^ values are presented in Figure S5.

| $RR_A (N_A)$ | DEXSeq | rMATS | SUPPA | MAJIQ-nw | MAJIQ-lw | MAJIQ-lw-relaxed |
| --- | --- | --- | --- | --- | --- | --- |
| **100% (80M)** | 0.5755 (>30000) | 0.734 (2595) | 0.822 (1790) | 0.864 (1234) | 0.861 (1232) | 0.780 (2111) |
| **50% (40M)** | 0.517 (28191) | 0.742 (1422) | 0.822 (1230) | 0.867 (340) | 0.873 (339) | 0.791 (1170) |
| **25% (20M)** | 0.485 (20349) | 0.760 (860) | 0.818 (852) | 0.885 (122) | 0.900 (120) | 0.799 (643) |

Table 1: **(a)** $RR_A$ and $N_A$ for subsampling of cerebellum and liver samples from [22] to 100% (80 million reads), 50% (40 million reads), and 25% (20 million reads) of their original coverage, as reported by DEXSeq, rMATS, SUPPA, MAJIQ-nw, MAJIQ-lw, and MAJIQ-lw with the relaxed filter as explained in the text.
| $R^2 (N)$ | rMATS | SUPPA | MAJIQ-nw | MAJIQ-lw | MAJIQ-lw-relaxed |
| --- | --- | --- | --- | --- | --- |
| **100% (80M)** | 0.959 (50) | 0.822 (37) | 0.967 (49) | 0.967 (49) | 0.967 (50) |
| **50% (40M)** | 0.936 (50) | 0.822 (37) | 0.904 (16) | 0.906 (16) | 0.941 (47) |
| **25% (20M)** | 0.952 (50) | 0.818 (37) | 0.914 (9) | 0.914 (9) | 0.924 (43) |

**Figure 4:**
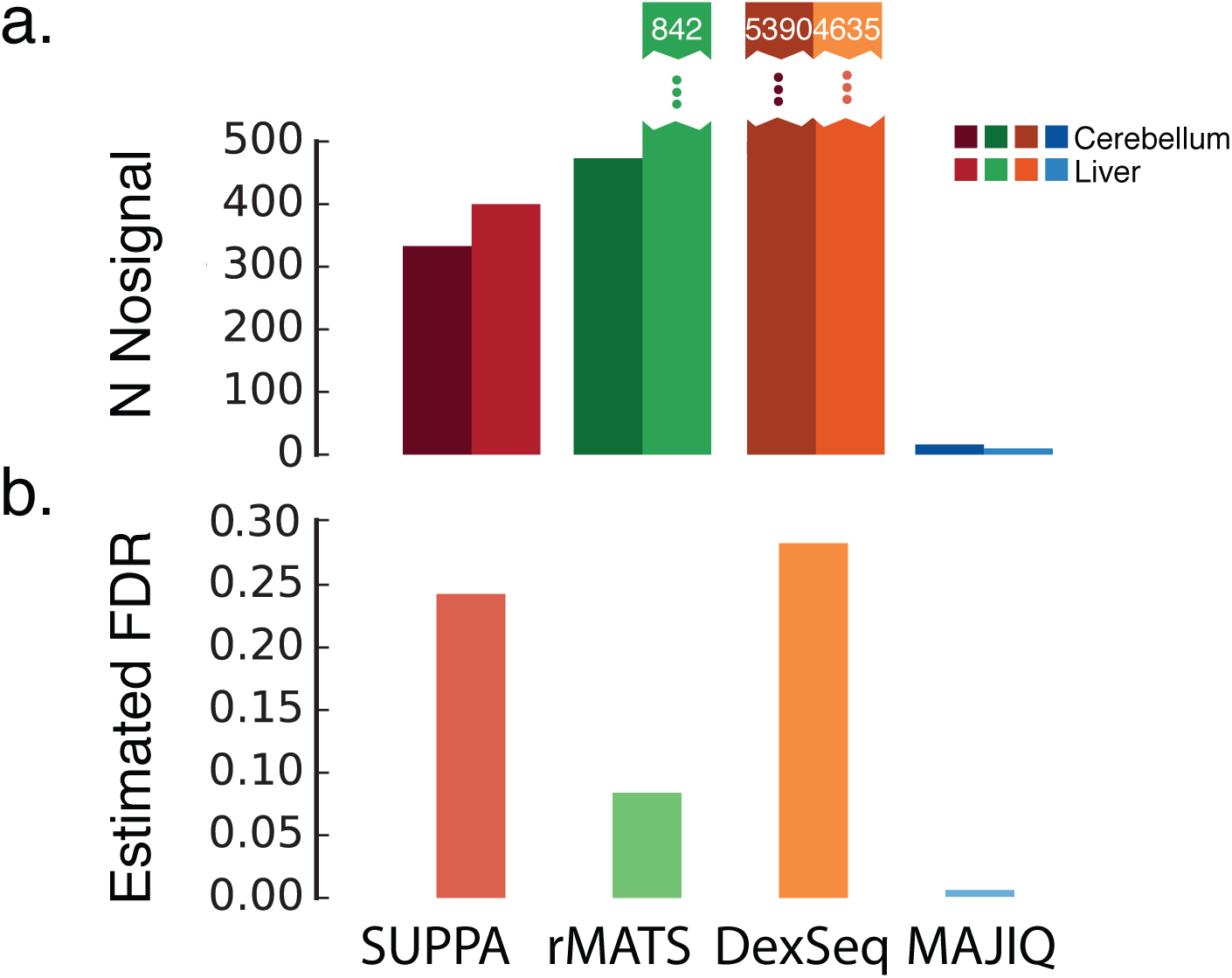
Evaluation of methods for “no-signal” experiment comparing groups of samples from the same condition (tissue type). **(a)** Number of detected events 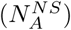. **(b)** Lower-bound estimate of false discovery rate (FDR) when comparing 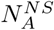 from (a) to *N*_*A*_ in Figure 3 (see Section 2.3).

Finally, we wanted to test how MAJIQ performs at different levels of read coverage. Unlike other methods, MAJIQ relies solely on junction-spanning reads and is therefore likely to suffer greatly in detection power if its default execution parameters are not adjusted accordingly. Specifically, we tested two different execution configurations. The first is MAJIQ’s default settings, which are particularly conservative and designed for high reproducibility assuming multiple replicates with an average coverage of approximately 60M and read length of 100 bases [22]. The second configuration, denoted “permissive”, drops the quantifiable filter restrictions to allow the detection of junctions in a single experiment with as few as 2 reads mapping across 2 or more positions on either side of the junction. We note that the other algorithms we analyzed do not have such coverage specific user adjustable parameters as these do no rely solely on junction spanning reads and as we show are less sensitive to changes in coverage.

By randomly sub-sampling the BAM files from [22], read coverage was varied from the original average of 80 million reads (denoted 100%) through 40 million (50%) to 20 million (25%). The reproducibility ratio and detection power of each method evaluated is reported in Figure 1a, and correlations with RT-PCR quantifications in Figure 1b. In general, all methods lose power to detect differentially-spliced events as coverage drops. As expected, MAJIQ run with default parameters is the most sensitive to sparse coverage, with detected events dropping 91%. When run with the relaxed parameters, MAJIQ loses 70%, similar to rMATS (67%) and more then SUPPA (53%) and DEXSeq (*≥* 34%). However, MAJIQ’s conservative settings pay off in consistent high reproducibility of up to 90%. In comparison, MAJIQ’s relaxed settings maintain low FDR (data not shown), but reproducibility is approximately 10% lower than the conservative settings. When evaluating the effect of sparse read coverage using the 50 RT-PCR experiments (Figure 1b), we find a similar picture. It is worth noting that these 50 events were designed by [19] to cover the wide range of read coverage levels across splicing events. In line with the observations from the reproducibility analysis, both the default and the relaxed settings maintain high *R*^2^ and compare favorably to other methods, but the default settings loses 82% of the events with an associated 6.3% drop in *R*^2^.

## 4 Discussion

In this paper, we developed a new model to automatically detect and downweight outliers in RNA-Seq datasets with replicates for splicing analysis. The problem of detecting outliers in batches of biological replicates has not received much attention in the literature as researchers are likely to simply discard samples before publication based on some heuristic such as PCA [5]. Such a heuristic may in turn reflect unconscious bias or cause good samples to be lost. Instead, we proposed a model for what constitutes an outlier sample in splicing analysis with a matching software that implements it. Users can run MAJIQ to evaluate possible outliers regardless of which pipeline they use for subsequent analysis.

In addition, we incorporated the outlier weighing scheme into the MAJIQ model either at the global, per-sample, level (MAJIQ-gw) or locally at the LSV level (MAJIQ-lw). Using both synthetic data perturbation and real data, we demonstrated that both MAJIQ-gw and MAJIQ-lw can maintain high levels of reproducibility when confronted with outliers. Furthermore, even when a sample is an outlier, using the local weight model (MAJIQ-nw) can help gain detection power by retaining LSVs which are “well-behaved”. In addition to sensitivity to outliers, we showed that low read coverage can also affect MAJIQ’s detection power as it relies on junction spanning reads. However, by adjusting MAJIQ’s default parameters, the new MAJIQ-lw was able to maintain high detection power with a moderate price in terms of reproducibility and correlation to RT-PCR, comparing favorably to other algorithms. Given that there is natural trade-off between detection power and reproducibility, we recommend users decide between default and permissive parameters based on their needs and dataset specifics.

While the new version of MAJIQ compared favorably to other algorithms in the experiments surveyed here, users should keep in mind that different datasets may suffer from different types of noise or biases, which may be different from the ones tested here. It is therefore advisable for potential users to always test DS algorithms using the evaluation criteria introduced here, including reproducibility plots, no-signal groups, and RT-PCR. To aid such evaluations, we made all the scripts used for this work available at majiq.biociphers.org.

Another important point to consider is that the methods tested here differ greatly in the set of features they offer. MAJIQ is the only one that offers the ability to detect complex splicing variations involving more than two alternative junctions, coupled with interactive visualization and genome browser connectivity. It is also capable of supplementing a given transcriptome annotation with reliable *de-novo* junctions detected in the RNA-Seq data, making it less dependent on the exact annotation. While useful for even normal tissues [19], this feature is particularly relevant for disease studies, cases where uncharacteristic splicing is expected, and for species with poorly-annotated transcriptomes. Notably, the latest version of rMATS, used in this work, offers to include *de-novo* junctions, but only adds those to annotated exons and requires the junctions to be at a predefined distance from those exons. In contrast, MAJIQ is able to detect completely novel exons and junctions. This ability of MAJIQ does come with a price of algorithm complexity and, consequently, execution time. While we did not perform detailed benchmarking, MAJIQ was much faster than rMATS and DEXSeq. However, SUPPA was much faster than all the other methods, as it assumes a known transcriptome and uses fast pseudoalignment algorithms such as SALMON [14] to quantify each transcript’s abundance. These assumptions may have deleterious effects on performance and might be at least partially responsible for the higher rate of false positives we observed for SUPPA. Finally, we note that other algorithms may have other specific desirable features. While including all of those is beyond the scope of this work users should consider such features as we well. For example, VAST-TOOLS [8] offers sensitive detection of micro-exons, and DiffSplice [7] offers detection of alternative splicing modules (ASM) in each gene splice graph.

There are several important directions in which this work can be extended. First, MAJIQ can be further improved both in terms of memory consumption and running time. While we were able to process over 100 samples with the current implementation on machines with 64GB of memory, parsing several hundreds or thousands or samples is currently not feasible. Furthermore, all the algorithms compared here were designed for datasets with small sets of biological replicates. Large heterogeneous datasets, such as those created in cancer studies, are likely to benefit from different statistical models. Finally, MAJIQ’s improved quantifications can be used to subsequently derive new models for splicing codes and splicing predictions given genetic variations [3, 4, 21]. Such improvements form a promising path for future algorithm development.

## Acknowledgements

We would like to thank the anonymous ISMB 2017 Reviewer for helpful suggestions on an earlier version of this work, as well as Matthew R. Gazzara and Anupama Jha from the BioCiphers Lab for helpful comments and suggestions.

### Funding

This work has been supported by R01 AG046544 to YB.

---

**Figure S1:**
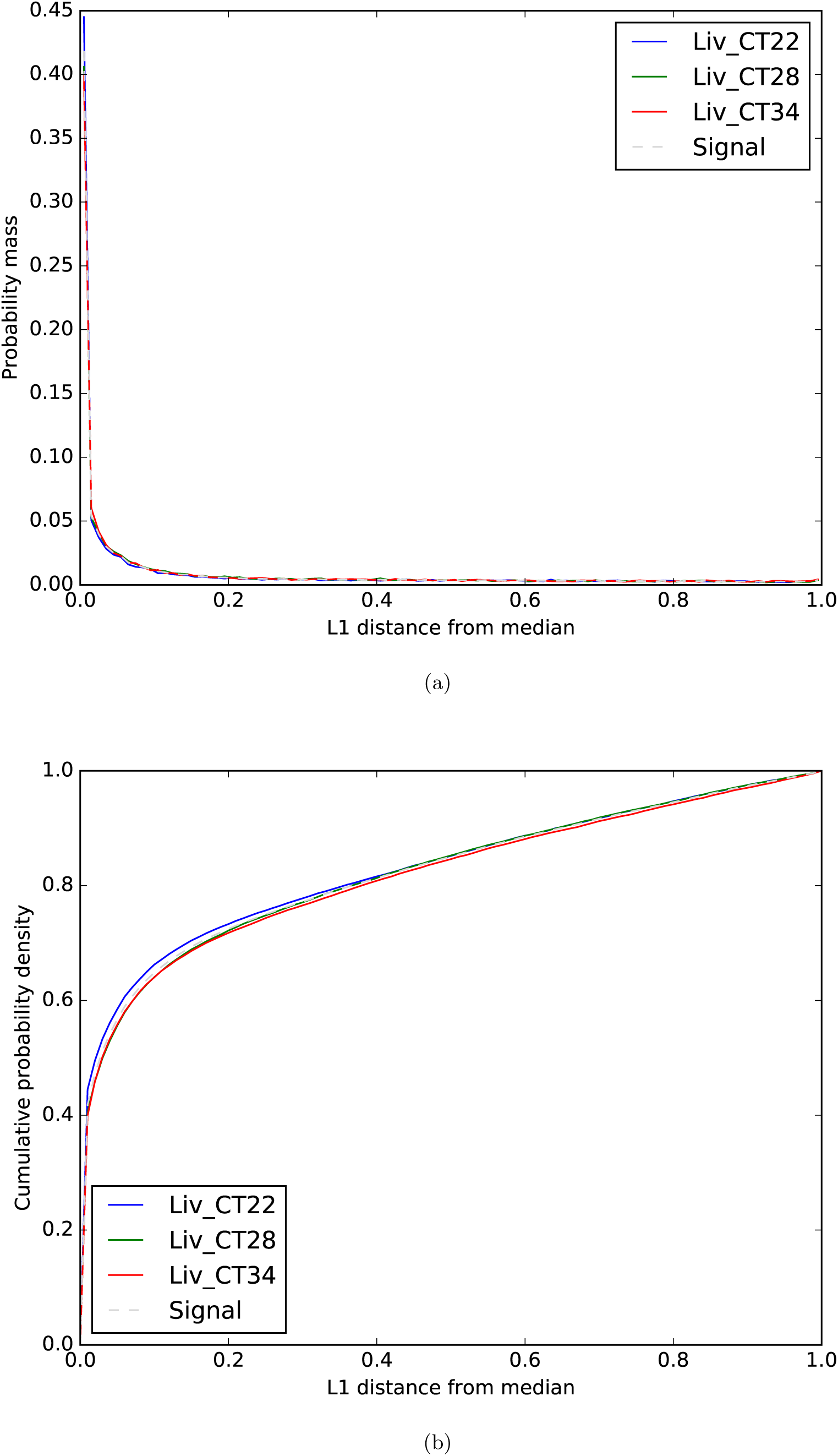
**(a)** Distribution of L1 divergences for three liver timepoints, to be used as a negative control for Figure 1. **(b)** Cumulative probability density for the same experiment. 90% of LSVs fall below the default threshold *τ* = 0.75.

**Figure S2:**
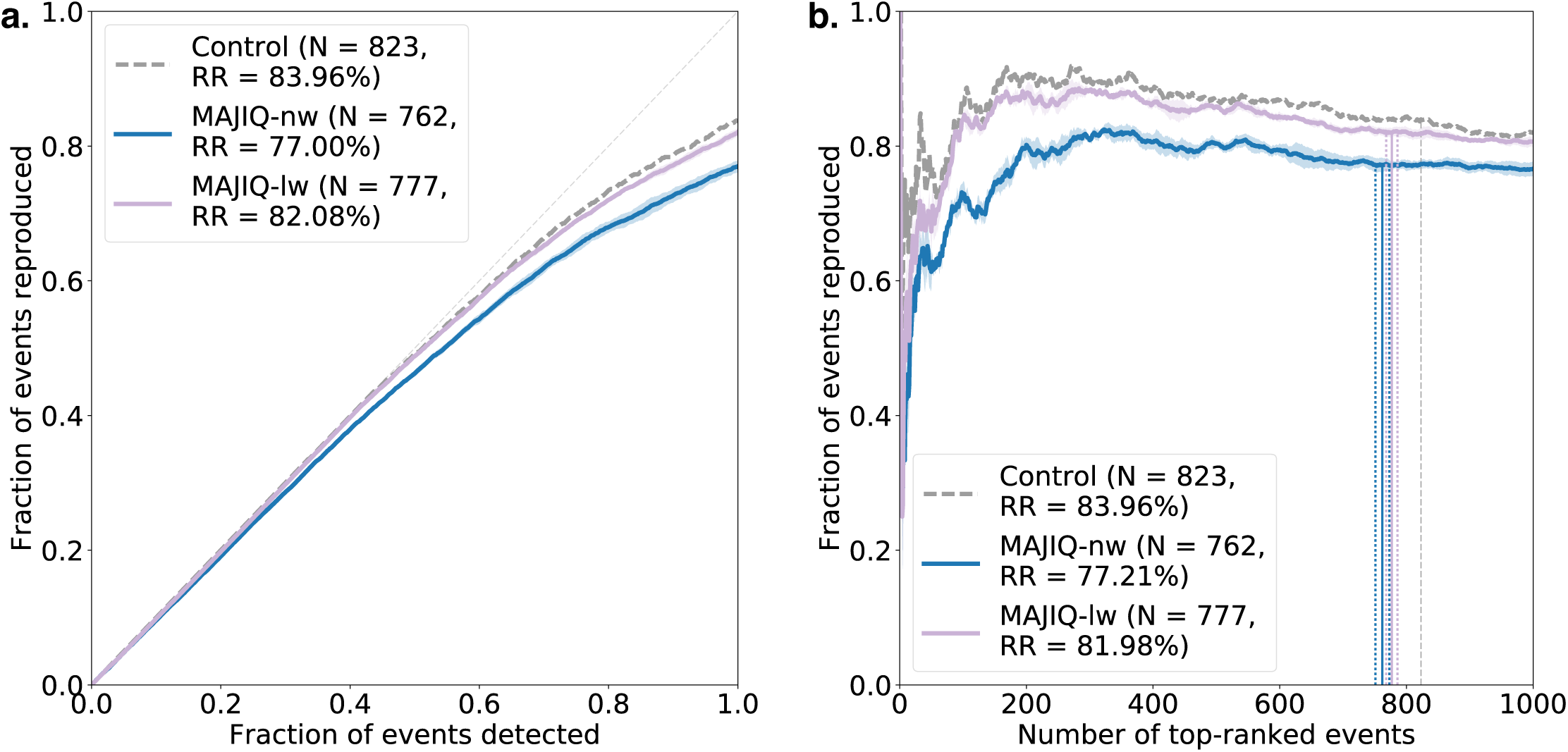
Side-by-side comparison of the two reproducibility plots. **(a)** The plot used in [19], where *RR*(*n*) is the fraction of the top *n* events in set 1 which also lie within the top *N*_*A*_ events in set 2. **(b)** The plot presented in Section 2.3, where *RR*(*n*) is the fraction of the top *n* events in set 1 which also lie within the top *n* events in set 2. Note that *RR*(*N*_*A*_), denoted by the dashed lines, is the same here as in (a).

**Figure S3:**
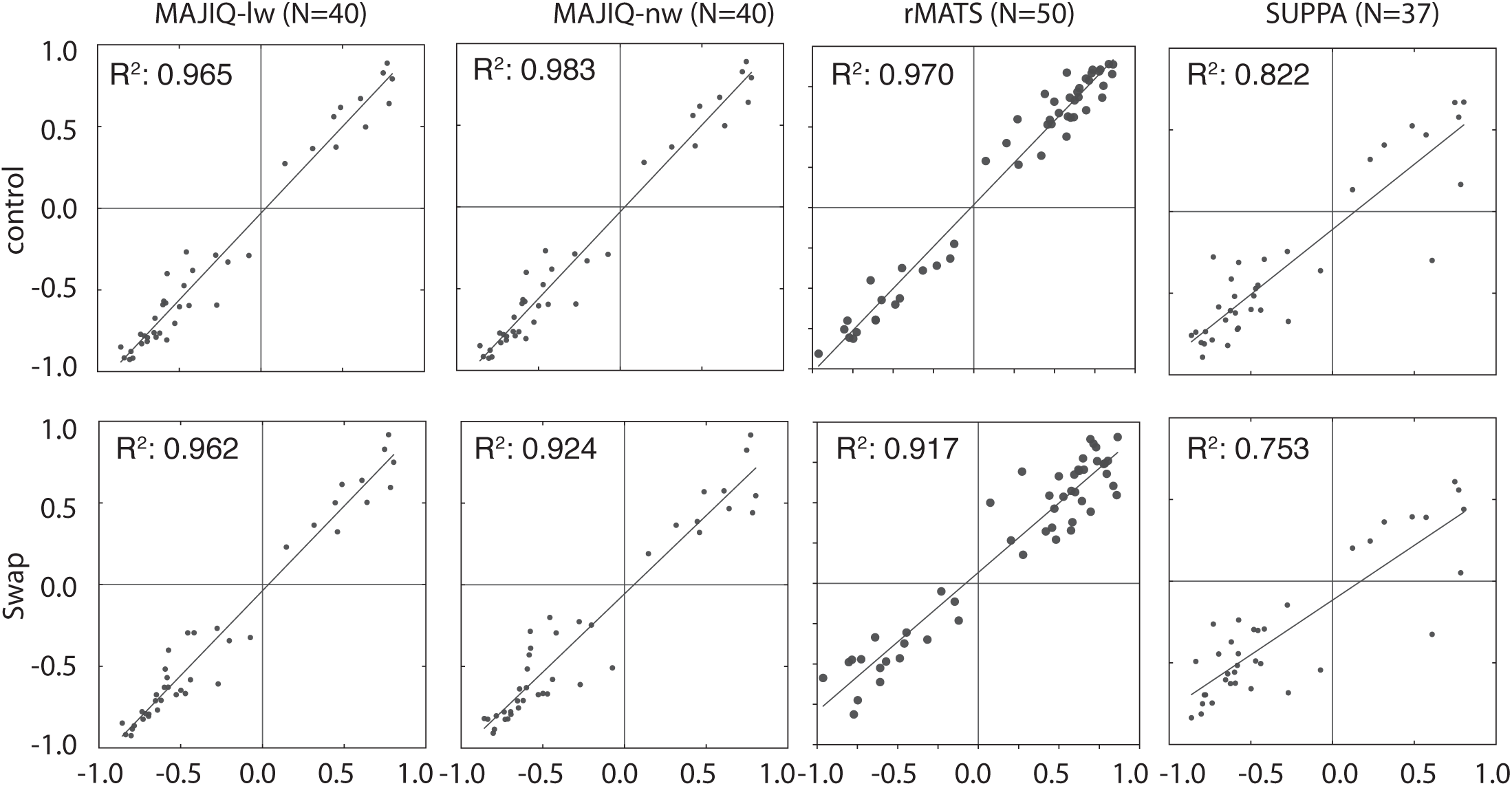
Scatterplots comparing RT-PCR ΔΨ quantifications with each model’s predictions with and without a swap. **Control** Cerebellum vs. liver, no swap. **Swap** Cerebellum vs. liver, with one cerebellum swapped out for a muscle sample. Each column show a different tool.

**Figure S4:**
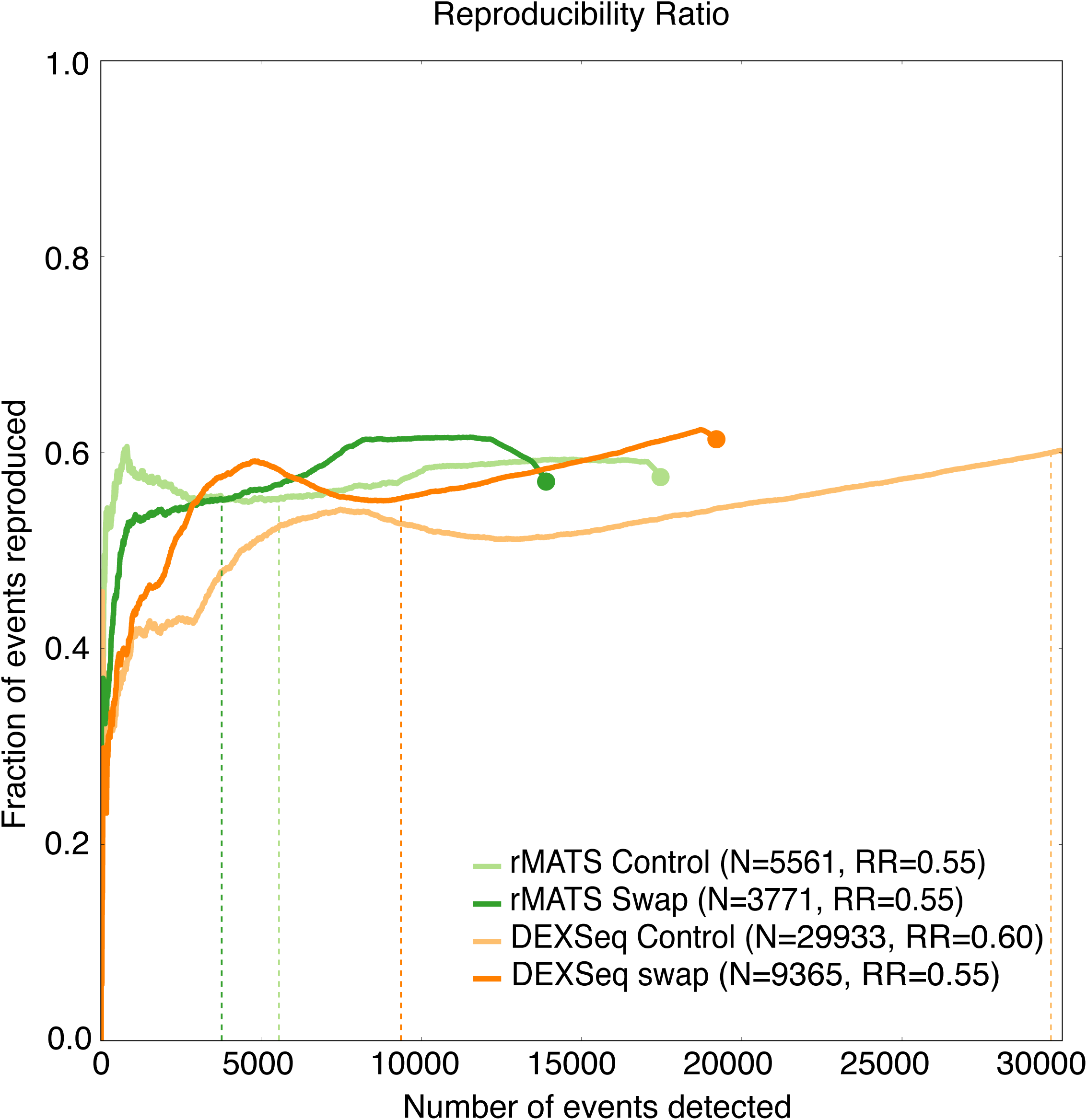
Extended RR plot for DEXSeq and rMATS showing the large number of splicing events flagged as significantly changing in the replicate swap experiment (Figure 3). Events beyond the 30000th call were ignored.

**Figure S5:**
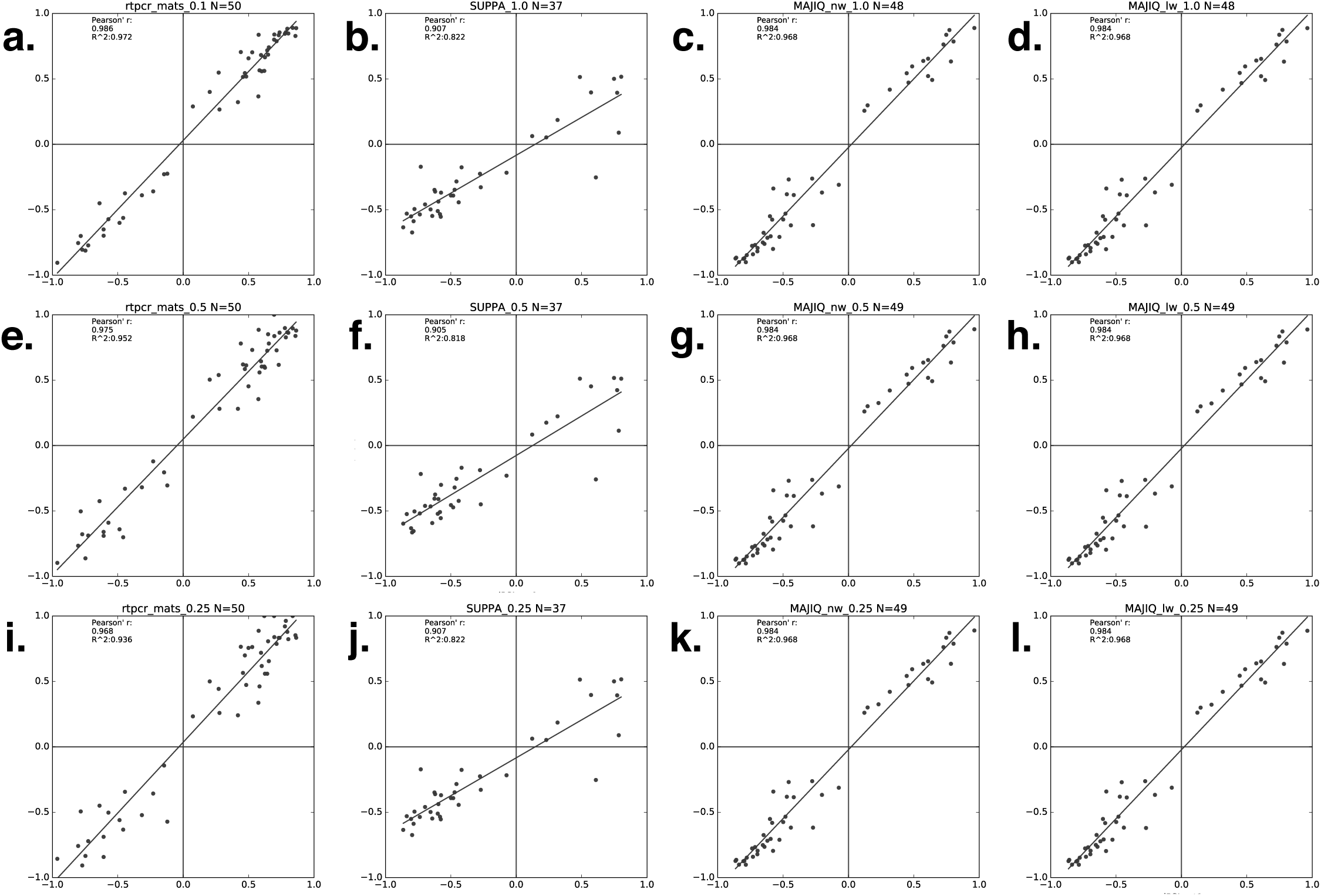
RT-PCR correlations for subsampling ratios 100% (a-d), 50% (e-h), and 25% (i-l) for rMATS (a,e,i), SUPPA (b,f,j), MAJIQ-nw (c,g,k), and MAJIQ-lw (d,h,l) as presented in Figure 1.

